# The IL-12– and IL-23–Dependent NK-Cell Response is Essential for Protective Immunity Against Secondary *Toxoplasma Gondii* Infection

**DOI:** 10.1101/547455

**Authors:** Daria L. Ivanova, Tiffany M. Mundhenke, Jason P. Gigley

## Abstract

Natural Killer (NK) cells can develop memory-like features and contribute to long-term immunity in mice and humans. NK cells are critical for protection against acute *T. gondii* infection. However, whether they contribute to long-term immunity in response to this parasite is unknown. We used a vaccine challenge model of parasite infection to address this question and to define the mechanism by which NK cells are activated during secondary parasite infection. We found NK cells were required for control of secondary infection. NK cells increased in number at the infection site, became cytotoxic and produced IFNγ. Adoptive transfer and NK-cell fate mapping revealed that *T. gondii*–experienced NK cells were not intrinsically different from naïve NK cells with respect to their long-term persistence and ability to protect. Thus, they did not develop memory-like characteristics. Instead, a cell-extrinsic mechanism may control protective NK-cell responses during secondary infection. To test the involvement of a cell-extrinsic mechanism, we used anti-IL-12p70 and IL-12p35^-/-^ mice and found that the secondary NK-cell response was not fully dependent on IL-12. IL-23 depletion with anti-IL-23p19 *in vivo* significantly reduced the secondary NK-cell response, suggesting that both IL-12 and IL-23 were involved. Anti-IL-12p40 treatment, which blocks both IL-12 and IL-23, eliminated the protective secondary NK-cell response, supporting this hypothesis. Our results define a previously unknown protective role for NK cells during secondary *T. gondii* infection that is dependent on IL-12 and IL-23.

## Introduction

Accumulating studies show that NK cells can acquire features of adaptive immune cells and develop immunological memory in response to certain stimuli (1). These memory-like NK cells provide a qualitatively and quantitatively greater response to secondary challenge and are intrinsically different from naïve cells. Antigen-specific memory NK cells are generated after encounters with haptens (2) and viruses, such as murine cytomegalovirus (MCMV) and human cytomegalovirus (HCMV) (3–5). *In vitro* and *in vivo* stimulation with certain cytokines, such as IL-12, IL-18 and IL-15, leads to the formation of memory-like features in NK cells that are epigenetically and functionally distinct from naïve cells (6–8). Both antigen specific and cytokine-activated memory-like NK cells are generated after MCMV infection *in vivo* (9). Whether NK cells develop memory-like characteristics in response to eukaryotic agents has yet to be found.

*T. gondii* is a food-borne intracellular parasitic protozoan that causes the disease toxoplasmosis. The parasite is present in one-third of the human population worldwide and is a significant health concern for immunocompromised individuals (10–13). At present, there is no vaccine or drug available to prevent or completely cure toxoplasmosis in humans (14, 15). NK cells are involved in innate immunity during acute *T. gondii* infection and are critical for early protection (16, 17). They mediate protection *via* IFNγ that is secreted in response to IL-12 provided by innate immune cells such as dendritic cells and macrophages (17, 18). NK-cell IFNγ also facilitates the differentiation of monocytes into inflammatory macrophages and monocyte-derived dendritic cells that then serve as the main source of IL-12 (19). In response to systemic IL-12 production during acute infection, bone marrow NK cells produce IFNγ and prime monocytes for regulatory function (20). NK cells also trigger an adaptive immune cell response to *T. gondii.* In the absence of CD4+ T cells, IFNγ produced by NK cells promotes CD8+ T-cell activation (21). In the absence of CD8+ T cells, NK-cell IFNγ contributes to the activation of CD4+ T cells (22). In addition to cytokine production, NK cells produce perforin and granzymes in response to the parasite and its subcellular components (23–26).

NK cells are clearly important for early control of *T. gondii* infection, yet their role in long-term immunity has not been addressed. This is clinically important to understand because there currently is no vaccine that elicits sterilizing immunity to the parasite (15, 27). A vaccine targeting the stimulation of NK cells in addition to CD8+ T cells could therefore be more beneficial long-term. In addition, *T. gondii* infection causes health complications in immunodeficient patients, many of whom are T-cell deficient (e.g., HIV patients) (11). Discovering new ways to utilize NK cells could be therapeutically beneficial for these patients.

In this study, we aimed to find whether NK cells contribute to long-term immunity against *T. gondii* in a vaccine challenge setting. We also investigated whether NK cells developed memory-like features in response to this vaccination. Lastly, we tested mechanisms involved in the activation of NK cells during secondary challenge. We demonstrate that NK cells are critical for reducing parasite burdens after lethal challenge. *T. gondii* infection induces a similar Th1 cytokine milieu as compared to MCMV, however, unlike memory-like NK cells generated by viral infection and cytokine stimulation (3, 9, 28), *T. gondii*–experienced NK cells did not intrinsically develop memory-like traits. This highlights for the first time that, NK cells are required for control of *T. gondii* reinfection, but are activated in this capacity by cell extrinsic mechanisms. Our exploration of the mechanisms involved in this secondary NK cell response revealed that their response to reinfection is dependent upon both IL-12p70 and IL-23. Our results reveal a novel role for NK cells during secondary *T. gondii* challenge infection in the presence of memory T cells that is dependent on IL-12 family cytokine stimulation.

## Materials and methods

### Mice

C57BL/6 (B6), CBA, B6.129S7-*Rag1^tm1Mom^* (*Rag1^-/-^*, Rag1 knockout [KO]), B6.129S1-*Il12a^tm1Jm^* (IL-12p35 KO), B6.129S 1*-Il12^tm1Jm^*/J (IL-12p40 KO) and B6.129X1-*Gt(ROSA)26Sor^tm1(EYFP)Cos^* (R26R-EYFP) mice were purchased from The Jackson Laboratory. B10;B6-*Rag2^tm1Fwa^ Il2rg^tm1Wjl^* (*Rag2^-/-^γc^-/-^*) mice were purchased from TACONIC. Transgenic NKp46-CreERT2 mice were kindly provided by Dr. Lewis Lanier (UCSF, CA) and crossed onto ROSA26R-EYFP mice to create inducible reporter mice for fate mapping. All animals were housed under specific pathogen-free conditions at the University of Wyoming Animal Facility.

### Ethics Statement

This study was carried out in strict accordance following the recommendations in the Guide for the Care and Use of Laboratory Animals of the National Institutes of Health. The University of Wyoming Institutional Animal Care and Use Committee (IACUC) (PHS/NIH/OLAW assurance number: A3216-01) approved all animal protocols.

### T. gondii *parasites and infection*

Tachyzoites of RH and RHΔ*cps1-1* (CPS) (kindly provided by Dr. David Bzik, Dartmouth College, NH) were cultured by serial passage in human fetal lung fibroblast (MRC5, ATCC) cell monolayers in complete DMEM (supplemented with 0.2 mM uracil for CPS strain). For mouse infections, parasites were purified by filtration through a 3.0-μm filter (Merck Millipore Ltd.) and washed with phosphate-buffered saline (PBS). Mice were infected intraperitoneally (i.p.) with 1 × 10^3^ or 1 × 10^6^ RH tachyzoites or 1 × 10^6^ CPS tachyzoites. The brains of CBA mice 5 wk after ME49 infection were used as a source of ME49 cysts. Mice were infected i.p. or i.g. (intragastrically) with 10 or 100 ME49 cysts.

### Cell depletion and fate mapping

To deplete NK cells, B6 mice were treated i.p. with 200 μg of anti-NK1.1 (PK136, Bio X Cell) 1 d before infection (d −1), on the day of infection (d 0) and every other day after infection for a maximum of 3 wk. To deplete CD8+ or CD4+ T cells, mice were treated i.p. with 200 μg of anti-CD8 T (2.43, Bio X Cell) or anti-CD4 T (GK1.5, Bio X Cell), respectively on d −1 and 0. To neutralize IL-12 family cytokines, mice were treated i.p. with 200 μg of anti-IL-12p70 (R2-9A5, Bio X Cell), anti-IL-12p40 (C17.8, Bio X Cell) and anti-IL-23p19 (MMp19B2, Biolegend) on d −1, 0 and 2. To block DNAM-1, mice were treated i.p. with 100 μg of anti-DNAM1 (480.1, Biolegend) on d −1 and 0. To induce yellow fluorescent protein (YFP) expression on NKp46+ cells, NKp46-CreERT2^+/−^ × R26R-EYFP^+/+^ mice were treated i.g. with 3.75 mg (males) and 2.5 mg (females) of tamoxifen (Sigma) for five consecutive days, beginning on the day of CPS immunization.

### Flow cytometry

Single-cell suspensions of peritoneal exudate cells (PECs), spleen and bone marrow (BM) were harvested from mice. Cells were then plated at 0.5−1.5 × 10^6^ cells/well. For surface staining, cells were washed twice with PBS and stained for viability in PBS using Fixable Live/Dead Aqua (Invitrogen) for 30 min. After the cells were washed with PBS, surface staining was performed using antibodies diluted in stain wash buffer (2% fetal bovine serum in PBS and [2 mM] EDTA) for 25 min on ice. For functional NK-cell assays, cells were stimulated for 4 h with plate bound anti-NK1.1 in the presence of 1× protein transport inhibitor cocktail (PTIC) containing Brefeldin A/Monensin (eBioscience, Thermo Fisher Scientific) and anti-CD107a (eBio1D4B, eBioscience, Thermo Fisher Scientific) in complete Iscove’s DMEM medium (Corning). After fixable live/dead and surface staining, the cells were fixed and permeabilized for 1 h on ice in Fixation/Permeabilization solution (BD Bioscience), followed by intracellular staining in 1× permeabilization wash buffer (BD Bioscience) with anti-IFNγ (XMG1.2, eBioscience, Thermo Fisher Scientific) for 40 min. Additional cells were surface stained with antibodies from Biolegend against CD3 (17A2), CD49b (DX5), NKp46 (29A1.4), NK1.1, CD4 (RM4-5), CD8b (YTS156.7.7), CD11b (M1/70), CD27 (LG.3A10), KLRG1 (2F1/KLRG1) and DNAM-1 (10E5); antibodies from eBioscience (Thermo Fisher Scientific) against Ly49I (YLI-90), Ly49H (3D10), CD94 (18d3) and CD107a (1D4B) and an antibody from BD Biosciences against Ly49D (4E5). The cells were washed twice with 1× PBS and resuspended in 1× PBS and analyzed using Guava easyCyte 12HT flow cytometer (Millipore) and FlowJo software (Tree Star).

### Adoptive transfer

Spleens were harvested from B6 mice 5 wk after immunization with CPS, from mice 5 wk after an initial immunization followed by a second immunization at wk 2 (2×CPS) and from naïve mice, and NK cells were purified by negative selection using EasySep Mouse NK Cell Isolation Kit (STEMCELL TECHNOLOGIES). Normalized NK-cell numbers (1–3 × 10^6^) were injected intravenously (i.v.) into *Rag2^-/-^γc^-/-^* or into naïve B6 mice. Recipient *Rag2^-/-^γc^-/-^* mice were infected with 1 × 10^3^ tach. RH i.p. or 10 ME49 cysts i.g. for the survival assessment. Recipient B6 mice were infected with 1 × 10^5^ tach. RH i.p. or 10 ME49 cysts i.p., and organs were harvested at 4 d after RH and 5 wk after ME49 infection for the parasite burden assessment.

### Parasite burden assessment with real-time PCR

DNA was extracted from the entire PEC and spleen sample harvested from infected mice using a DNeasy Blood & Tissue Kit (Qiagen). Parasite DNA from 600 ng of PEC DNA and 800 ng of splenic tissue DNA was amplified using primers specific for the *T. gondii* B1 gene (forward primer 5’-GGAACTGCATCCGTTCATG-3’ and reverse primer 5’-TCTTTAAAGCGTTCGTGGTC-3’) at 10 pmol of each per reaction (Integrated DNA Technologies) by real-time fluorogenic PCR using SsoAdvanced Universal IT SYBR Green SMx (BIO-RAD) on a CFX Connect Real-Time System cycler (BIO-RAD). Parasite equivalents were determined by extrapolation from a standard curve amplified from purified RH parasite DNA.

### Statistical analysis

Statistical analysis was performed using Prism 7.0d (GraphPad) and Microsoft Excel 2011. Significant differences were calculated using either unpaired Student’s t-test with Welch’s correction or analysis of variance (ANOVA). The log-rank (Mantel-Cox) test was used to evaluate survival rate. Data is presented in graphs as the mean± standard deviation (SD). Significance is denoted as follows: ns, not significant (p > 0.05) or significant with a p-value less than 0.05.

## Results

### NK cells are required for protection during secondary *T. gondii* infection

NK cells have been shown in several infection models to develop memory-like characteristics, including the ability to contribute to protection against secondary challenge of infection (3, 29–31). NK cells are critical for protection during acute *T. gondii* infection (16); however, whether they can also contribute to secondary infection by this parasite in the presence of T-cell memory is not known. A previously published report suggested that NK1.1+ and ASGM1+ cells helped provide immunity against *T. gondii* reinfection in the absence of CD8+ T cells (32). Therefore, we used a vaccine challenge model to test whether NK cells also helped provide immunity against *T. gondii* in the presence of memory T cells. For the primary infection, B6 mice were infected i.p. with tachyzoites of live attenuated uracil auxotroph strain CPS. These parasites are able to invade cells *in vivo* but do not replicate in the absence of uracil and get cleared within a week (33). Infection with this vaccine strain induces localized inflammation and leads to the generation of memory CD8+ T cells that protect mice against lethal reinfections regardless of the infecting strain or route of infection (34, 35). Five to 6 wk after CPS immunization, mice were challenged i.p. with lethal doses of highly virulent type I RH tachyzoites. To determine if NK cells were important during this secondary challenge, NK cells were depleted using anti-NK1.1 upon reinfection, and parasite burdens were measured by semi-quantitative real-time PCR for the *T. gondii*–specific B1 gene. As shown in Figure 1A, parasite burdens were significantly higher in NK cell–depleted mice as compared with the controls. To investigate the importance of NK cells during secondary challenge with type II ME49 parasites and the long-term infection, CPS-immunized mice were orally challenged with the type II ME49 strain and treated with anti-NK1.1 for 3 wk. Mouse cyst burden and survival were assayed at 5 wk after infection. The cyst counts in the brain clearly showed elevated cyst numbers in anti-NK1.1-treated mice as compared with undepleted controls (Fig. 1B). These data indicate that NK cells play a protective role in secondary infections after vaccination and in the presence of T-cell memory during type I (RH) and type II (ME49) *T. gondii* infection.

**FIGURE 1.**
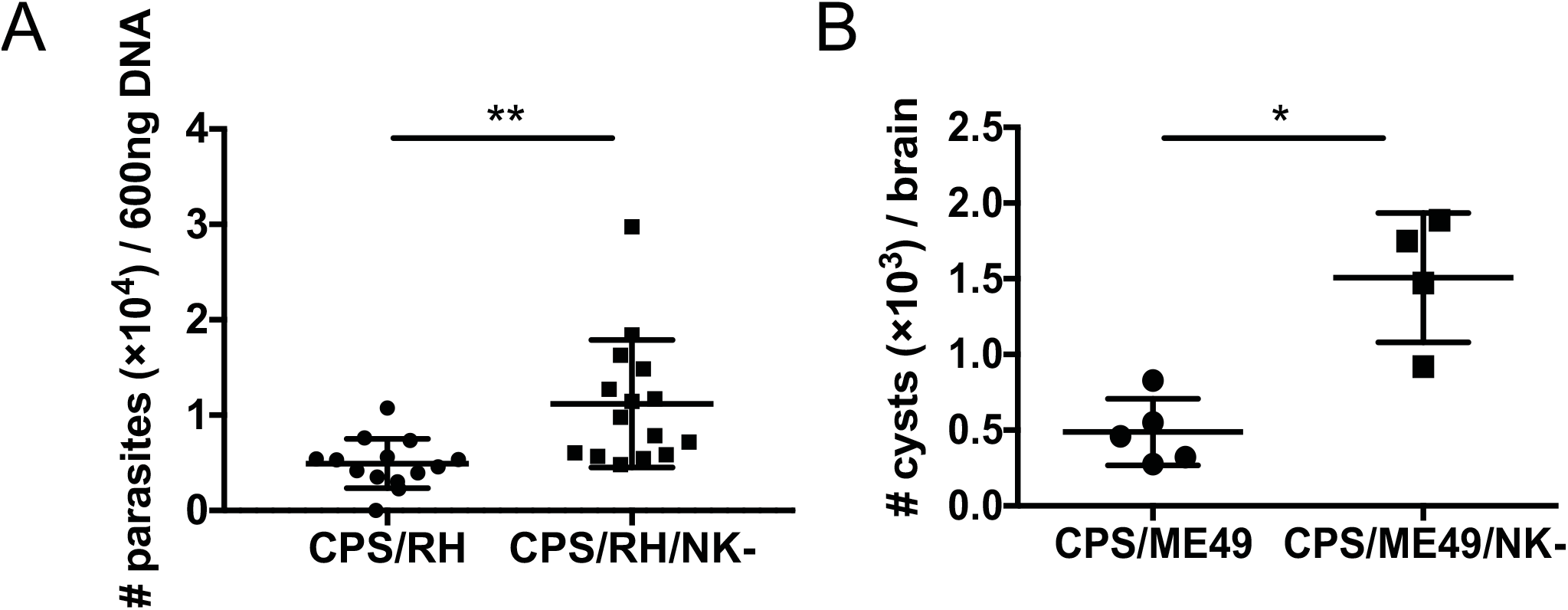
NK cells are required for protection during secondary *T. gondii* infection. (**A** and **B**) B6 mice were infected i.p. with 1 × 10^6^ CPS. Five to 6 wk later, mice were infected i.p. with **(A)** 1 × 10^6^ RH tachyzoites or **(B)** 100 brain ME49 cysts and were treated with anti-NK1.1 (PK136) or were untreated. **(A)** Two days after RH infection, the number of parasites was compared among PEC DNA (600 ng/sample) based on semiquantitative real-time PCR for the *T. gondii* B1. Concatenated data from four independent experiments, n = 3 or 4 mice/group. **(B)** *T. gondii* cysts were quantified in the brains of B6 mice 5 wk after ME49 infection. Representative data from one of two independent experiments, n = 3–5 mice/group. Data are the mean ± SD with individual data points. Unpaired Student’s t-test with Welch’s correction, *p < 0.05, **p < 0.01.

### NK cells become activated during an adaptive recall response

We showed in Figure 1 that, after vaccination and in the presence of memory T cells, NK cells contributed to the control of a challenge infection. How the NK cells were responding to the secondary infection was not, however, known. Therefore, we next determined whether NK cells were activated during secondary challenge and defined how they responded. We measured NK-cell frequency, number and functionality in the peritoneum, the site of infection, and in the spleen by flow cytometry. B6 mice were infected with CPS i.p. and 5 to 6 wk later were challenged i.p. with type I RH parasites. NK1.1+CD3− lymphocytes, which included NK cells and other innate lymphoid cells (ILCs) that express the NK1.1 receptor, increased in frequency and number at the infection site as early as 1 d after reinfection, remained there through d 5 and gradually declined by d 10 (Fig. 2A, 2B and 2C). Over the course of the secondary infection, NK1.1+CD3− cells constituted a large proportion of the lymphocytes with frequencies and numbers comparable to those of CD8+ T cells (Fig. 2B, 2C). To differentiate between NK cells and other NK1.1+ ILCs, NK cells were further defined as CD49b+NKp46+ cells (gated on CD3− live lymphocytes) (Fig. 2D). Only CD49b+NKp46+ cells increased in frequency and number at the site of reinfection, suggesting that ILC1 (CD49b−NKp46+) did not mount a secondary response (Fig. 2D, 2E). NK cells did not increase in the spleen after challenge, suggesting that the secondary NK-cell response was localized to the site of infection (Supplemental Fig. 1A, 1B). In the peritoneum, the NK-cell increase in frequency and number after secondary infection was similar to that which occurred during primary infection (Supplemental Fig. 1C).

**FIGURE 2.**
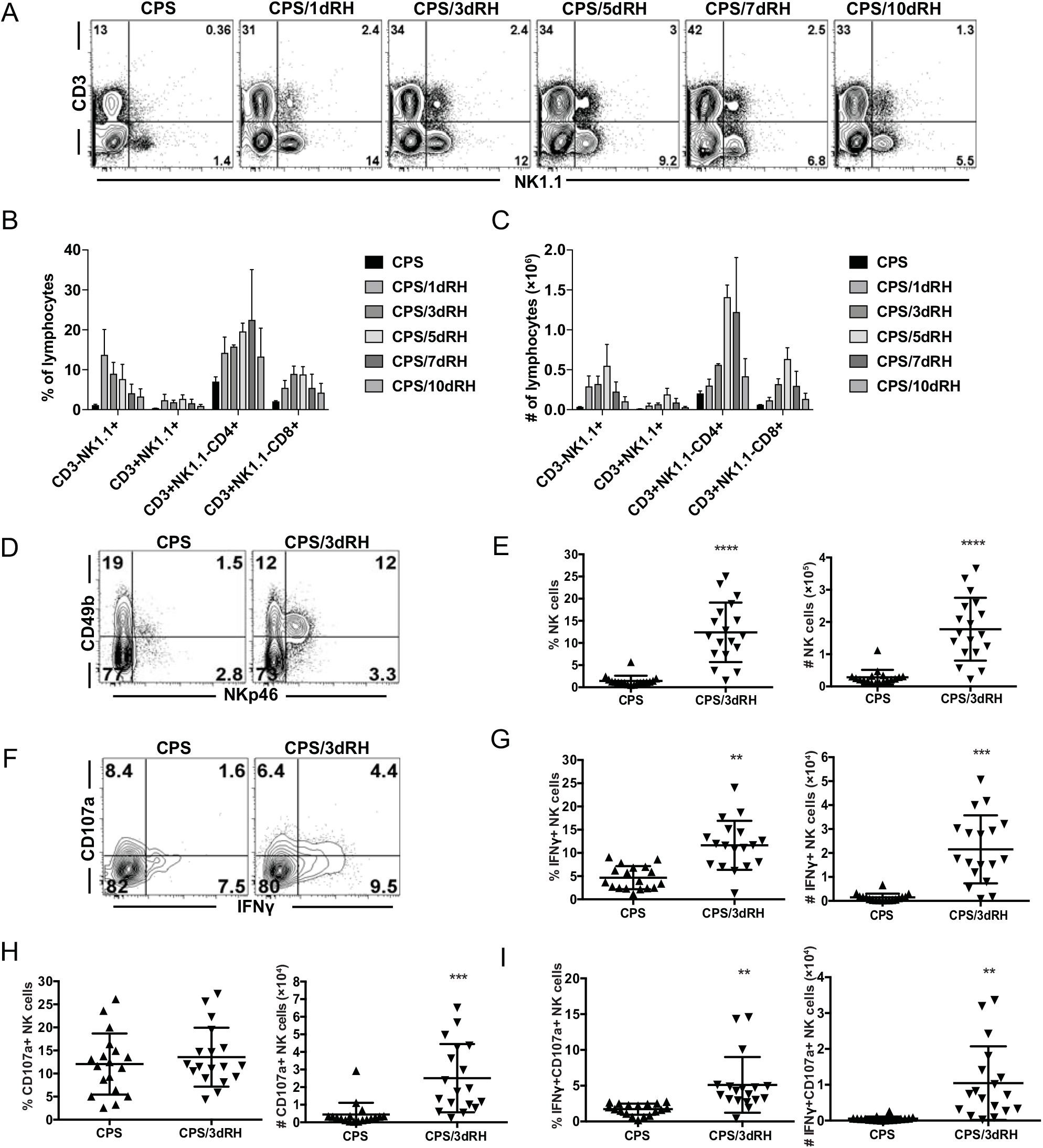
NK cells become activated during an adaptive recall response. **(A-I)** B6 mice were infected i.p. with 1 × 10^6^ CPS tachyzoites and were then reinfected i.p. with 1 × 10^3^ RH tachyzoites 5–6 wk later. PECs were analyzed by flow cytometry at d 1, 3, 5, 7 and 10 (**A, B** and **C**) or d 3 **(D-I)** after RH infection. **(A)** Representative contour plots of the cells expressing NK1.1 and CD3 in live lymphocytes. **(B)** The percentages and **(C) numbers** per PEC of CD3−NK1.1+ (ILCs), CD3+NK1.1+ (NK T cells), CD3+NK1.1−CD4+ (CD4+ T cells) and CD3+NK1.1−CD8b+ (CD8+ T cells) among live lymphocytes. Data from one of two independent experiments, n = 3 or 4 mice/group. Data are the mean ± SD. **(D, E)** The percentages and **(E)** numbers of NK cells (CD49b+NKp46+) among CD3− live lymphocytes. (FI) The percentages and numbers of IFNγ+, CD107a+ and IFNγ+CD107a+ NK cells. **(D-I)** Concatenated data from four experiments, n = 3 or 4 mice/group. Data are the mean ± SD. **p < 0. 01, ***p < 0.001, ****p < 0.0001, one-way ANOVA.

The main function provided by NK cells during acute *T. gondii* infection is IFNγ production, which is critical for protection (16, 36). To assess how NK cells were responding, we measured their IFNγ production. NK cells produced IFNγ during secondary infection, and the frequency and number of IFNγ+ NK cells increased at the site of infection (Fig. 2F, 2G and Supplemental Fig. 1D). In addition to the cytokine production, NK cells develop a cytotoxic response after stimulation with *T. gondii* parasites and their subcellular components (23, 25). We measured NK-cell cytotoxicity during secondary challenge using the surrogate marker CD107a (37). The frequency of cytotoxic CD107a+ NK cells did not significantly change, whereas their absolute cell numbers increased (Fig. 2F, 2H and Supplemental Fig. 1D). NK cells also developed a polyfunctional response (CD107a+IFNγ+), which was significantly increased during secondary challenge (Fig. 2F, 2I and Supplemental Fig. 1D). These findings indicate that NK cells become rapidly activated during the response to secondary *T. gondii* infection to provide effector functions constituting a substantial portion of the total lymphocytes during a recall response.

### Multiple NK-cell subpopulations are activated during secondary *T. gondii* infection

Inflammatory cytokines and/or activating receptor engagement with cognate ligands can activate NK cells (38, 39). After MCMV infection, both cytokine-activated and ligand-driven NK-cell responses occur (9). During some viral infections, specific NK-cell subpopulations, defined by their receptor expression, dominate the response (40, 41). During acute *T. gondii* infection, we and others have not observed a dominant responding NK-cell population, suggesting that NK cells are driven by a cytokine-dependent process, rather than by a process that depends on activating receptor ligands (22, 42). To further define how NK cells were responding to secondary *T. gondii* infection, we measured whether a dominant NK-cell population developed in the context of immune memory to the parasite. Mice were immunized with CPS and 5 to 6 wk later were infected with type I RH. The expression of inhibitory receptors, including Ly49I and CD94, and activating receptors, including Ly49H and Ly49D, was assessed on the total NK-cell population as well as on IFNγ+ NK cells by flow cytometry. Total and IFNγ+ NK cells were similarly distributed within the subsets of lymphocytes most of which showed the expected activation in the RH-infected mice relative to the CPS-only mice, and there was no dominant population observed after reinfection (Fig. 3A, 3B). Interestingly, the fold increase in the CD94+ subset was higher than that of Ly49I+, Ly49H+ and Ly49D+ NK cells.

**FIGURE 3.**
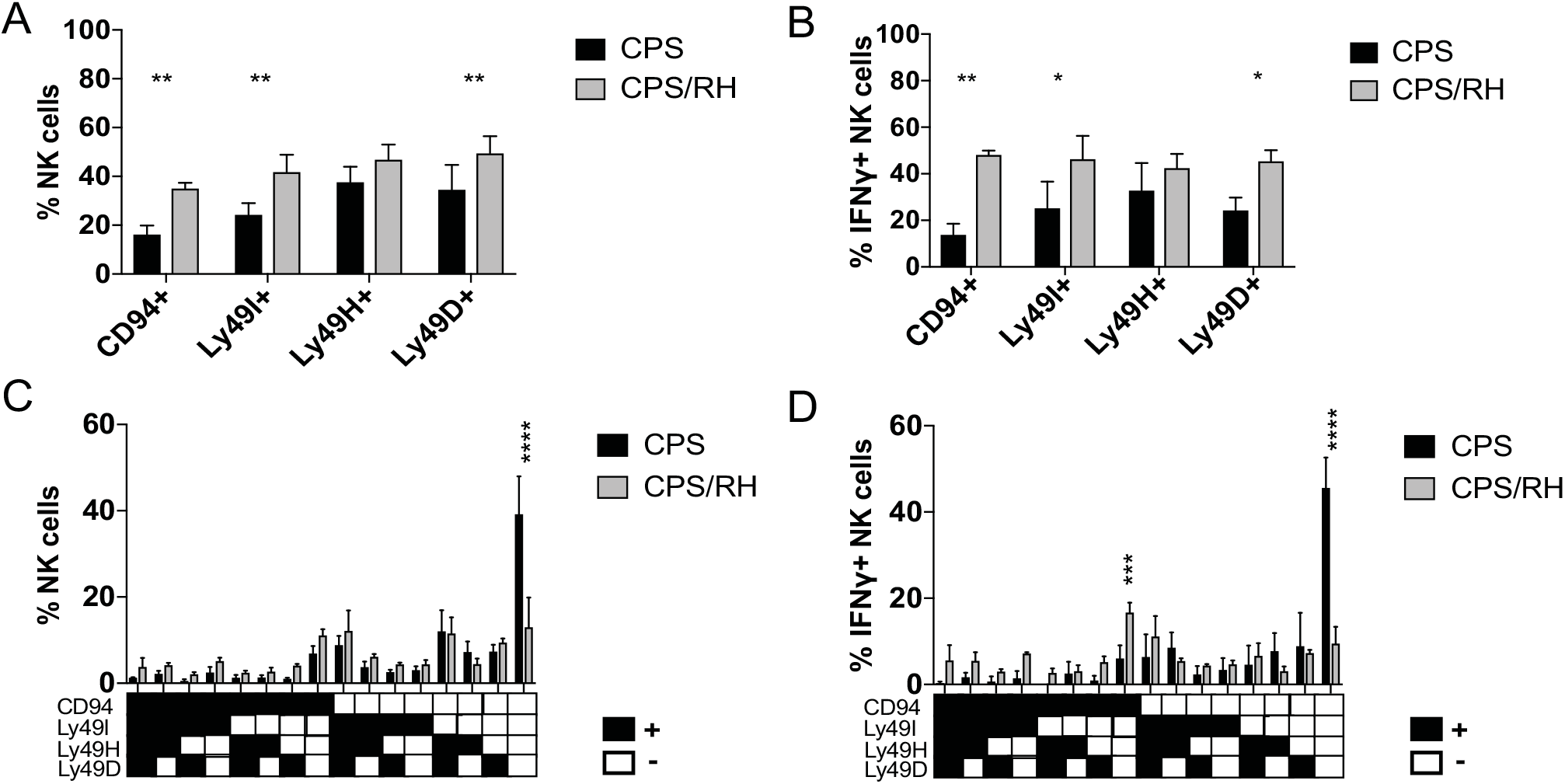
Multiple NK-cell subpopulations are activated during secondary *T. gondii* infection. **(A-D)** B6 mice were infected i.p. with 1 × 10^6^ CPS and 5–6 wk later were infected i.p. with 1 × 10^3^ RH tachyzoites. PECs were analyzed by flow cytometry at d 3 after RH infection. (A and B) The frequency of CD94+, Ly49I+, Ly49H+ and Ly49D+ cells within the total **(A)** and IFNγ+ **(B)** NK cells. (**C** and **D**) The frequency of NK cells expressing combinations of the receptors (CD94, Ly49I, Ly49H, Ly49D) within the total **(C)** and IFNγ+ (D) NK cells. Representative graphs from one of two independent experiments with n = 4 mice/group. Data are the mean ± SD. *p < 0.05, **p < 0.01, ***p < 0.001, ****p < 0.0001, one-way and two-way ANOVA.

As NK cells stochastically express an array of receptors, the dominant NK-cell population might express multiple receptors rather than a single receptor. We therefore next analyzed the phenotype of NK cells expressing combinations of NK-cell receptors in CPS-immunized mice compared with mice that were immunized with CPS and later challenged with RH 3 d before analysis. Among the total NK cells and IFNγ+ NK cells, we observed a significant decrease in the frequency of CD94−Ly49I−Ly49H−Ly49D−NK cells in the RH-challenged mice (Fig. 3C, 3D). This reduction of the receptor-negative NK cells was associated with an increase in NK cells that expressed multiple combinations of the receptors and that were widely distributed. Interestingly, the combinations that included CD94 receptor had the greatest increase in frequency (Fig. 3D). This included NK cells that were CD94+Ly49I−Ly49H−Ly49D−. Nevertheless, compared with viral infections, a distinct, dominant responding NK-cell population did not stand out during secondary *T. gondii* infection, similar to our observations during acute *T. gondii* infection.

### Activated NK cells during secondary *T. gondii* infection are mature

MCMV-primed Ly49H+ and alloantigen-primed Ly49D+ ligand-driven NK-cell subsets that develop intrinsic memory develop a mature phenotype marked by being CD11b+CD27−KLRG1^high^DNAM−1^low^ on their surface (3, 43, 44). Maturation of NK cells then seems to be critical for their ability to acquire features of adaptive immune cells. Therefore, measuring whether NK cell maturation was altered during secondary challenge after vaccination is important to better understand NK-cell biology in this setting. We next defined how vaccination and the presence of immune memory impacted NK-cell maturation after secondary *T. gondii* infection. As NK cells mature in the periphery, they progress from the least mature stage, R0 (CD27−CD11b−); to stage R1 (CD27+CD11b−); followed by stage R2 (CD27+CD11b+) and the terminally mature stage, R3 (CD27−CD11b+) (45). Mice were immunized with CPS as described above, and NK-cell maturation was measured at the site of reinfection based on expression of CD27 and CD11b. RH infection of CPS-vaccinated animals significantly elevated NK-cell maturation to stage R2 (Fig. 4A). This was even more evident when we measured NK cell maturation (CD27−CD11b+) in IFNγ+ NK-cell populations (Fig. 4B). Immature NK cells (CD27−CD11b−, R0) decreased, intermediately mature NK cells (CD27+CD11b+, R2) increased and other subsets (R1, CD27+CD11b+ and R3, CD27−CD11b+) did not change after secondary infection.

**FIGURE 4.**
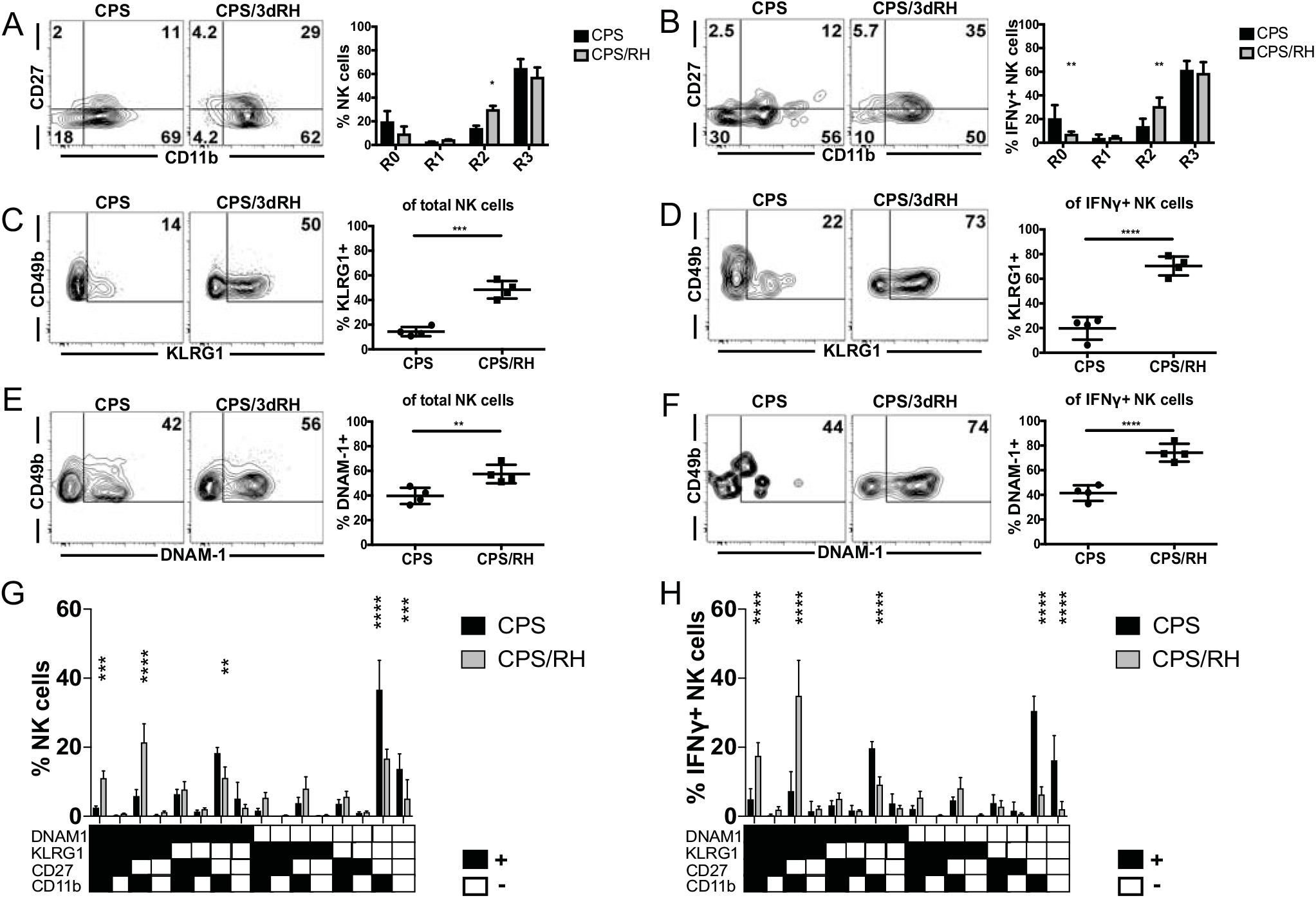
Activated NK cells during secondary *T. gondii* infection are mature. **(A-H)** B6 mice were infected i.p. with 1 × 10^6^ CPS and 5–6 wk later were infected i.p. with 1 × 10^3^ RH tachyzoites. PECs were analyzed by flow cytometry at d 3 after RH infection. (**A** and **B**) The frequency of R0 (CD27−CD11b−), R1 (CD27+CD11b−), R2 (CD27+CD11b+), R3 (CD27−CD11b+) NK cells within the total **(A)** and IFNγ+ **(B)** NK cells. (**C** and **D**) The frequency of KLRG1+ NK cells within total **(C)** and IFNγ+ **(D)** NK cells. (**E** and **F**) The frequency of DNAM-1+ NK cells within the total **(E)** and IFNγ+ **(F)** NK cells. (**G** and **H**) The frequency of NK cells expressing the combinations of the receptors (KLRG1, DNAM1, CD27, CD11b) within the total **(G)** and IFNγ+ **(H)** NK cells. Representative graphs from one of two independent experiments with n = 4 mice/group. Data are the mean ± SD. *p < 0.05, **p < 0.01, ***p < 0.001, ****p < 0.0001, one-way and two-way ANOVA.

In addition to conventional maturation markers, we also measured expression of the NK-cell activation marker KLRG1 and the costimulatory molecule DNAM-1. Adaptive NK cells have high KLRG1 and low DNAM-1 expression after MCMV and alloantigen stimulation (3, 43, 44). NK cells from CPS-immunized and naïve mice did not significantly differ in the expression of these markers (data not shown). There was a significant increase in KLRG1+ and DNAM-1+ subsets after reinfection (Fig. 4C-F). By measuring the expression of these markers in combination with each other, we found that NK cells of the memory cell phenotype (KLRG1^high^DNAM−1^low^CD11b+CD27−) did not significantly change and constituted a small portion of the total and IFNγ+ NK cells before and after reinfection (Fig. 4G, 4H). The population that increased and produced the most IFNγ was KLRG1+DNAM-1+CD11b+CD27−, followed by KLRG1+DNAM-1+CD11b+CD27+. IFNγ was mainly produced by mature DNAM-1+ rather than mature DNAM-1− NK cells (Fig. 4H). This suggested that DNAM-1 could be involved in NK-cell responses to secondary *T. gondii* infection.

We addressed this possibility by blocking DNAM-1 signaling *in vivo* using anti-DNAM-1 in CPS-immunized and RH-challenged mice. Anti-DNAM-1 treatment appeared to possibly deplete DNAM-1^high^ NK cells, however, treatment did not result in a reduction of total NK cell and IFNγ-producing NK cell frequency or number or frequency (Supplemental Fig. 2 A,B and C). Based on this result, treatment most likely did not deplete DNAM-1^high^ cells and simply blocked the receptor resulting in decreased staining *ex vivo* (Supplemental Fig. 2C). In summary, there does not appear to be a specific NK-cell subpopulation that develops in response to *T. gondii* vaccination.

### *T. gondii*–experienced and naïve NK cells are not intrinsically different

Accumulating studies indicate that NK cells can further differentiate after primary stimulation (cytokine stimulation, hapten exposure or viral infection) to acquire features of adaptive immune cells and develop memory-like abilities in a cell-intrinsic manner (2, 3, 31). However, the development of cell-intrinsic NK-cell memory-like features in response to eukaryotic pathogens has yet to be discovered. To address whether *T. gondii* infection-experienced NK cells were intrinsically different from naïve cells (without parasite experience) in their ability to protect against secondary infection, we used T and B cell-deficient *Rag1^-/-^* mice (46). One group of *Rag1^-/-^* mice were immunized with CPS parasites, whereas a second was not. Both groups were challenge infected with type I RH (i.p.) or type II ME49 cysts (i.p. and i.g.) 5 to 6 wk later, and their survival was monitored CPS-immunized *Rag1^-/-^* mice were not protected better than non-immunized controls (Fig. 5A-C). The administration of the parasites *via* different routes also did not affect this result (Fig. 5B, C).

**FIGURE 5.**
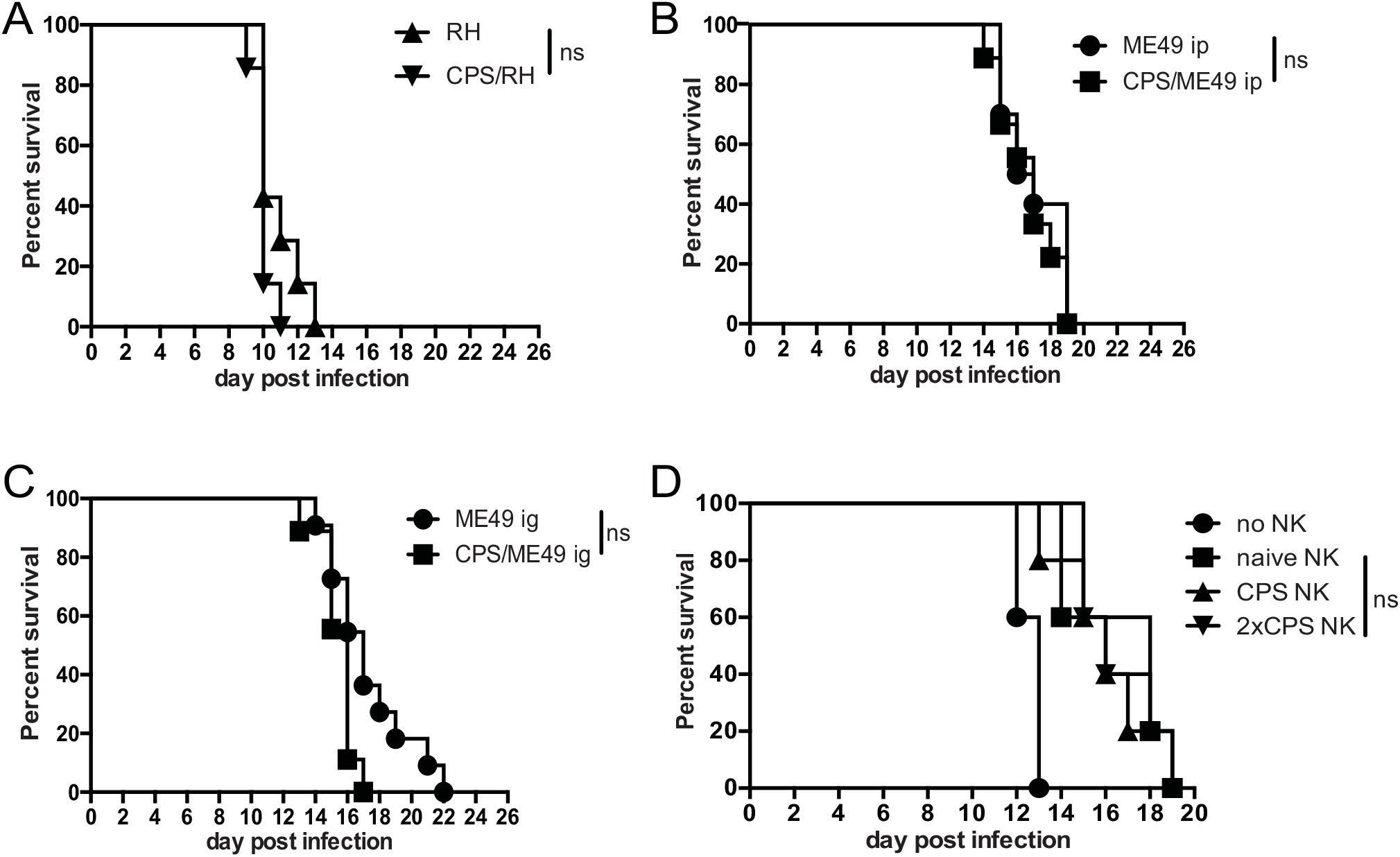
*T. gondii*–experienced and naïve NK cells are not intrinsically different with respect to their ability to protect against subsequent infection. **(A-C)** Survival after infection with 1 × 10^3^ RH tachyzoites i.p. **(A)** or with 10 brain ME49 cysts i.p. **(B)** or i.g. **(C)** was compared between *Rag1^-/-^* mice 5–6 wk after CPS immunization and non-immunized *Rag1*^-/-^ mice. Each graph represents concatenated data from two independent experiments, n = 3–6 mice/group. **(D)** NK cells were purified from spleens of naïve B6 mice (naive NK) and from mice 5 wk after CPS immunization (CPS NK) and from mice 5 wk after initial CPS immunization that was followed by a second round of immunization at wk 2 (2×CPS NK) B6 mice. Purified NK cells were then transferred i.v. into *Rag2^-/-^γc*^-/-^ mice (1.4 × 10^6^ NK cells/mouse). Recipient mouse survival after i.g. infection with 10 ME49 cysts was determined. The data represent one of two independent experiments, n = 5 mice per experiment. ns, not significant. The log-rank (Mantel-Cox) test was used to evaluate survival rates.

NK cells do not express RAG recombinase once they begin to develop (47). However, in *Rag1^-/-^* mice there is impaired NK-cell expansion and differentiation of memory-like characteristics after MCMV infection (48). One reason we may not have observed a survival difference between CPS-immunized *Rag1^-/-^* and non-immunized *Rag1^-/-^* mice could be due to the RAG1 deficiency and a consequent loss of the development of CPS immunization-induced NK cell-intrinsic memory-like features. Therefore, to further test whether cell-intrinsic differences existed between naïve and CPS-experienced NK cells, we performed NK-cell adoptive transfer into *Rag2^-/-^γc^-/-^* mice, which lack T, B and NK cells (49). Wild-type (WT) B6 animals were immunized with CPS parasites or were not immunized. Bulk NK cells were purified by negative selection from the spleens of 5-wk CPS-immunized mice or non-immunized age-matched controls. NK cells were transferred i.v. into the *Rag2^-/-^γc^-/-^* recipient mice. Those mice were then challenged with type II ME49 cysts, and their survival was measured. As in immunized *Rag1^-/-^* animals, *Rag2^-/-^γc^-/-^* recipients of CPS-experienced NK cells did not survive any longer than did recipients of naïve NK cells (Fig. 5D). This result indicated that *T. gondii*–experienced NK cells were not intrinsically different in their ability to protect immunodeficient mice as compared with naïve NK cells.

In *Rag2^-/-^γc^-/-^* mice, adoptively transferred NK cells undergo homeostatic proliferation, resulting in their spontaneous activation and production of IFNγ (50). This could have been why we could not detect a cell-intrinsic difference between *T. gondii*–experienced and naïve NK cells. Therefore, to limit homeostatic proliferation-associated cell activation, WT mice were used as recipients of CPS-experienced and naïve NK cells to see if parasite-experienced NK cells were more protective. Purified NK cells from the spleens of CPS-immunized or naïve B6 mice were transferred into WT recipients. Recipient animals were then challenged with either type I RH parasites or type II ME49 cysts. The parasite burdens were measured by real-time PCR for the parasite-specific B1 gene at 4 d after RH and by brain cyst counts 5 wk after ME49 infection. We did not observe any significant differences in parasite burdens between the recipients of CPS-immunized and naïve NK cells (Supplemental Fig. 3 A, 3B).

In addition to conveying more-efficient protection in response to secondary challenge with the same stimuli, memory lymphocytes develop the ability to persist longer than naïve cells (31). For the above adoptive transfer experiments, NK cells were purified from B6 mice 5 wk after immunization. These bulk NK cells contained cells that had experienced primary *T. gondii* infection as well as newly generated naïve cells. To differentiate between long-lived and naïve NK cells and to assess their ability to persist, the tamoxifen-inducible reporter strain NKp46-CreERT2 × Rosa26-YFP was used to fate map NK cells activated during immunization and not newly generated NK cells (9). To label and track whether NK cells were long-lived after *T. gondii* infection, reporter mice were immunized with CPS and at the same time treated with tamoxifen. Tamoxifen treatment was continued for five consecutive days after immunization, and mice were harvested 5 wk later. Control reporter mice that were not immunized with CPS were treated with tamoxifen for 5 d and harvested 5 wk later. The flow cytometry analysis revealed that the frequency and number of YFP+ NK cells, which represented long-lived persistent cells, was not different between CPS-immunized and naïve mice in the peritoneum (the site of infection) (Fig. 6A), spleen (periphery) (Fig. 6B) and the bone marrow (the site of NK-cell generation) (Fig. 6C). Interestingly, there was a trend toward a reduction in the frequency of YFP+ NK cells in immunized mice as compared with naïve mice. Taken together, these results indicate that *T. gondii*–experienced and naïve murine NK cells are not intrinsically different in their ability to protect against or persist after *T. gondii* infection.

**FIGURE 6.**
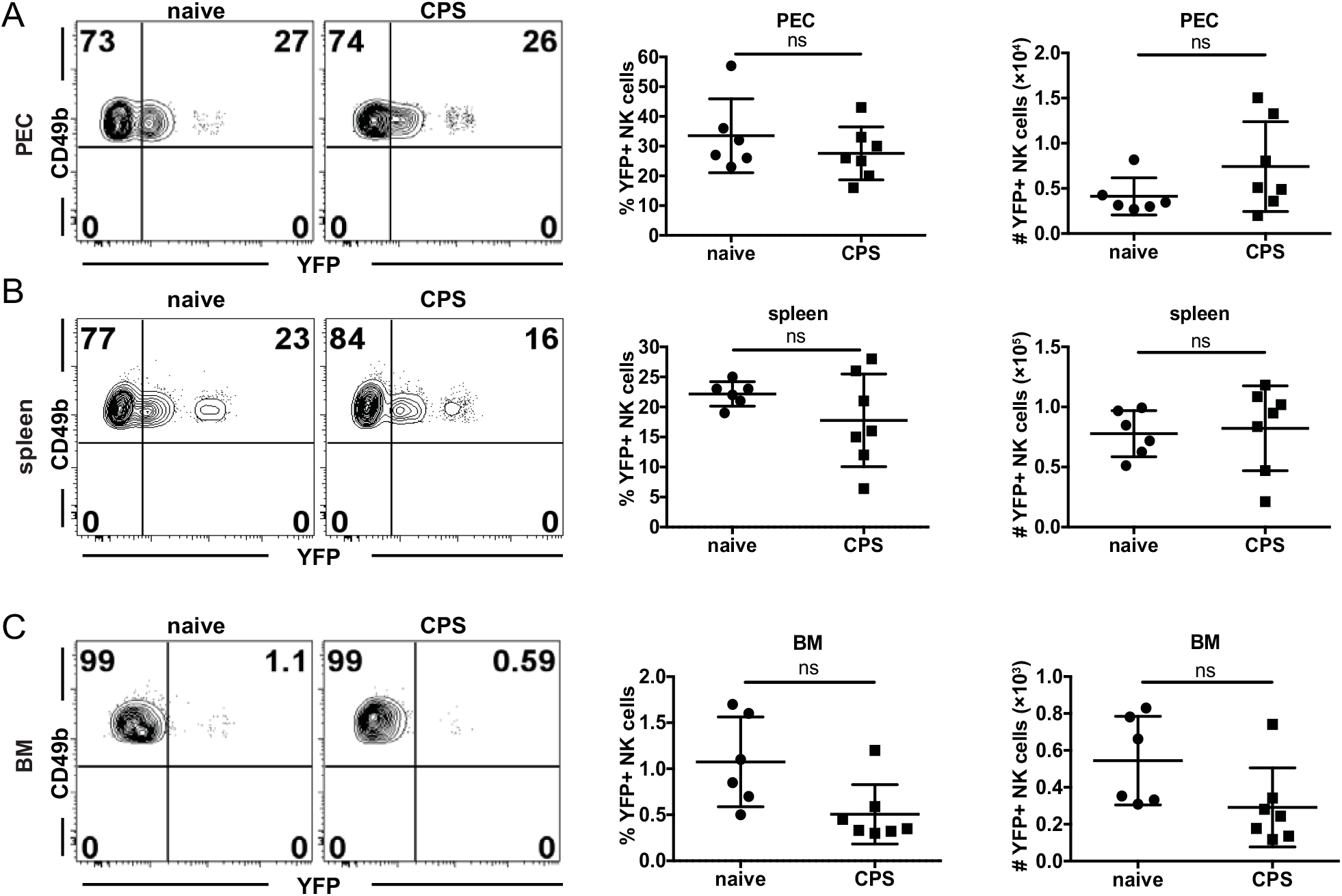
*T. gondii*–experienced and naïve NK cells are not intrinsically different in their ability to persist. **(A-C)** NKp46-CreERT2 × Rosa26-YFP mice were i.g. treated with tamoxifen for five consecutive days beginning on the day of CPS immunization. Five weeks after tamoxifen treatment, the frequency and number of YFP+ NK cells (gated as CD49b+NKp46+ within CD3− live lymphocytes) were compared between CPS-immunized and non-immunized mice by flow cytometry in the PEC **(A)**, spleen **(B)** and BM **(C)**. The data represent one of two experiments, n = 4–7 mice/group. Data are the mean ± SD. ns, not significant. Statistical analysis was performed using one-way ANOVA.

### NK cells get activated independently of T cells

Intrinsic NK-cell memory does not develop in all disease situations (51). NK cells can still contribute to adaptive immune recall responses even if they are not intrinsically different from naïve cells. Studies suggest that antigen-specific CD4+ T cells rapidly produce IL-2 and can activate NK cells (51–54). We found that NK cells responded to secondary *T. gondii* infection, but they were not intrinsically different from naïve NK cells. Therefore, we next examined whether memory CD4+ or CD8+ T cells were required for helping to activate NK cells in response to secondary *T. gondii* infection. To test the role of CD4+ and CD8+ T cells in NK-cell activation, mice were immunized with CPS and 5 to 6 wk later depleted of CD4+ or CD8+ T cells. Depleted and non-depleted animals were then challenged with RH, and NK-cell responses were measured at the site of infection. NK cells increased in frequency and number and produced IFNγ even when CD4+ or CD8+ T cells were absent (Fig. 7). Interestingly, after the treatment with anti-CD4, NK cells produced more IFNγ than non-treated mice. This could be explained by a concurrent depletion of CD4+ Treg cells by the antibody (55). Overall, the above result showed that, in contrast to other infection models in which NK-cell responses depended on antigen-specific T cells, CD4+ and CD8+ T cells were not required for NK-cell activation during secondary *T. gondii* infection.

**FIGURE 7.**
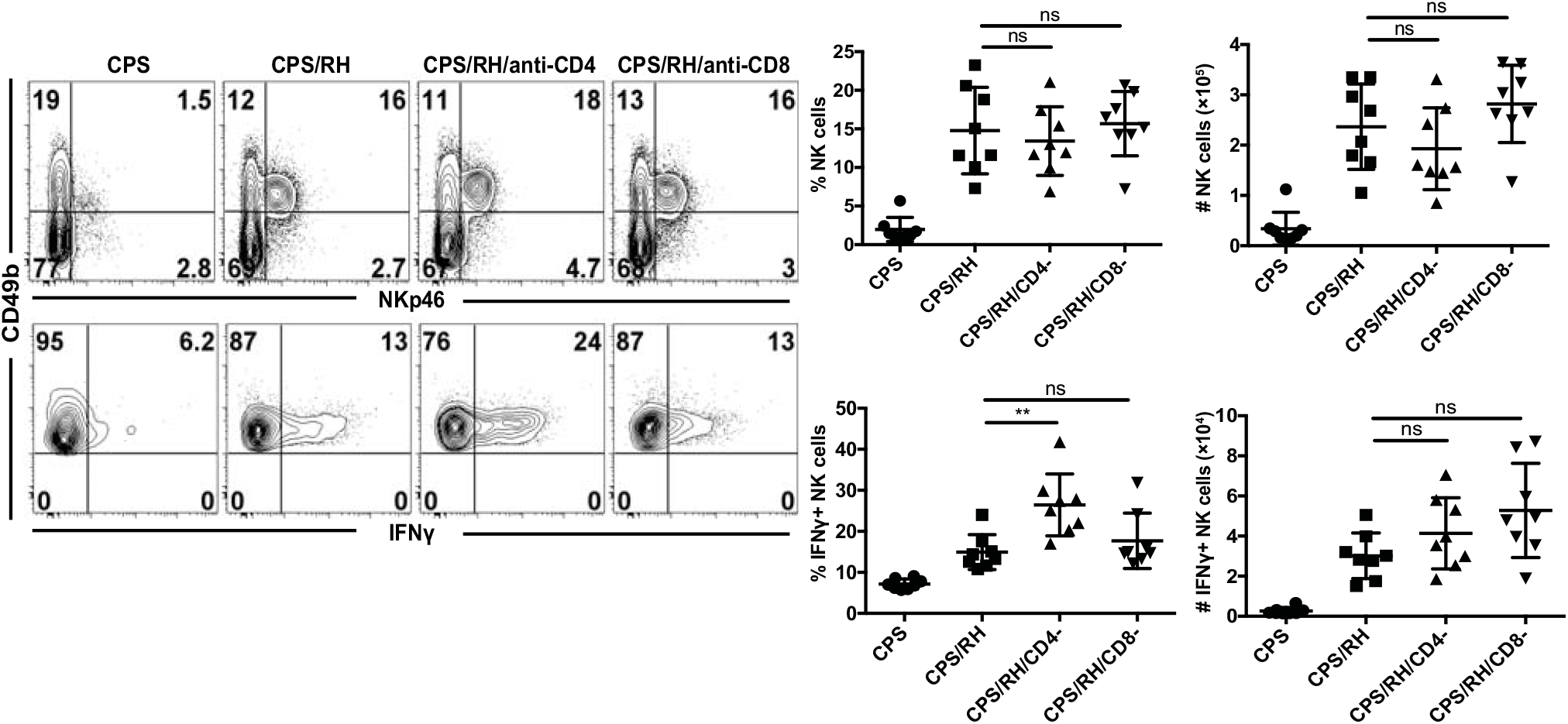
NK cells are activated independently of T cells. B6 mice were infected i.p. with 1 × 10^6^ CPS tachyzoites. Five to 6 weeks later, mice were infected i.p. with 1 × 10^3^ RH tachyzoites and treated with antibodies that deplete CD4+ or CD8+ T cells. Three days after RH infection, the frequency and number of NK cells (CD49b+NKp46+CD3− live lymphocytes) and their IFNγ production in PECs were measured by flow cytometry. Representative contour plots from one of two experiments. Concatenated graphs from two independent experiments, n = 4 mice/group. Data are the mean ± SD. ns, not significant; **p < 0.01, one-way ANOVA.

### IL-12 and IL-23 are required for the NK-cell response to secondary infection

*T. gondii* infection was one of the first systems in which the IL-12/IFNγ axis was demonstrated (17). More specifically, IL-12−dependent NK-cell IFNγ production was shown to protect against acute *T. gondii* infection (16–18). Thus, IL-12 could be required for NK-cell IFNγ production during secondary infection with the parasite in the presence of immune memory. To test whether IL-12 was involved in secondary NK-cell responses, CPS-immunized mice were treated with anti-IL-12p40 or were untreated and then were challenged with RH tachyzoites. The frequency and absolute number of total NK cells and activated IFNγ+ NK cells were measured 3 d later. The frequency of NK cells was not affected by anti-IL-12p40, but their numbers were significantly lower than in untreated mice (Fig. 8A). Consistent with the previously shown role of IL-12p40 during acute infection (17, 18), the absence of IL-12p40 during secondary infection led to a dramatic decrease in IFNγ production by NK cells. Both the frequency and number of IFNγ+ NK cells were comparable to non-challenged control mice. These data show that after secondary *T. gondii* infection, IL-12p40 is essential for NK-cell IFNγ production.

**FIGURE 8.**
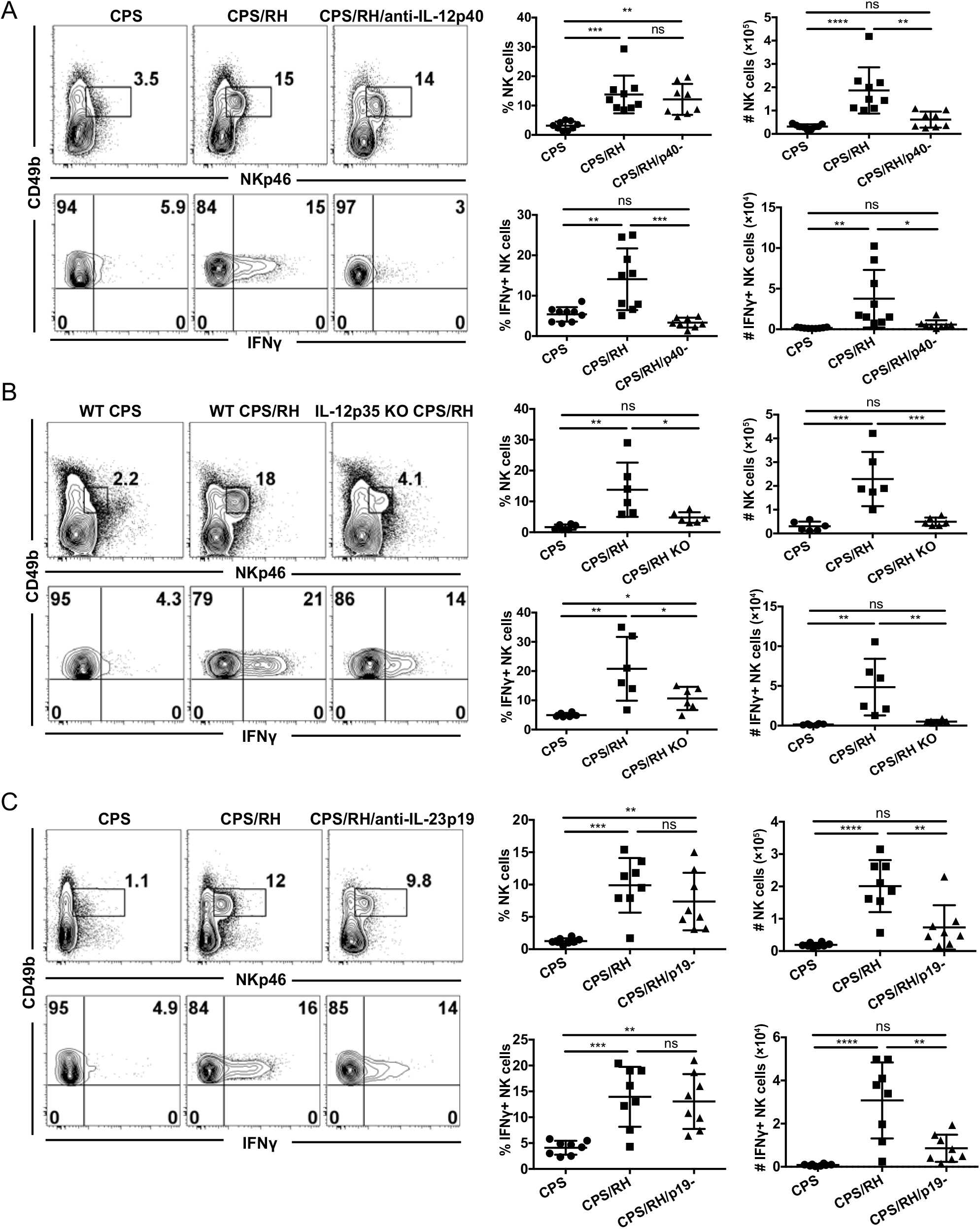
IL-12 and IL-23 are required for the NK-cell response to secondary *T. gondii* infection. **(A-C)** B6 (**A** and **C**) and IL-12p35 KO **(B)** mice were i.p. infected with 1 × 10^6^ CPS and 5–6 wk later were i.p. infected with 1 × 10^3^ RH tachyzoites. NK cells (CD49b+NKp46+CD3− lymphocytes) and their IFNγ production were analyzed in PECs at 3 d after RH infection by flow cytometry. **(A)** Mice were treated with anti-IL-12p40 blocking antibody during RH infection. Representative contour plots and graphs are shown from one of two independent experiments. Concatenated graphs from two independent experiments, n = 3–5 mice/group. **(B)** NK-cell frequency and IFNγ production in IL-12p35 KO mice. Representative contour plots from one of two independent experiments. Concatenated graphs from two independent experiments, n = 3 mice/group. **(C)** Mice were treated with anti-IL-23p19 during RH infection. Representative contour plots from one of two independent experiments. Concatenated graphs from two independent experiments, n = 4 mice/group. Data are the mean ± SD. ns, not significant; *p < 0.05, **p < 0.01, ***p < 0.001, ****p < 0.0001, ordinary one-way ANOVA.

In the majority of studies, the role of IL-12 in acute *T. gondii* infection has been assayed by measuring the p40 subunit of IL-12 (17, 18). However, p40 is a subunit of multiple cytokines and can be biologically active as a subunit of heterodimers IL-12p70 and IL-23 as well as mono- and homodimers (56–58). Therefore, we more specifically addressed the role of IL-12 in NK-cell activation during secondary infection by neutralizing IL-12p70 by antibody treatment during RH reinfection of CPS-immunized mice. Consistent with the anti-IL12p40 data, the blockade of IL-12p70 did not significantly affect NK-cell frequency, but NK-cell numbers were reduced (Supplemental Fig. 4). In contrast to anti-IL-12p40, anti-IL-12p70 treatment did not lead to a significant decrease in the frequency or number of IFNγ-producing NK cells. As this was an unexpected result, we further tested the role of IL-12p70 in NK-cell activation in IL-12p35 KO mice. Although T cells do not develop memory in the absence of IL-12 in IL-12p35 KO mice (59), we could use this mouse model because the NK-cell response to secondary *T. gondii* infection did not require T-cell help. Flow cytometry analysis revealed that both the frequency and number of total and IFNγ-producing NK cells in IL-12p35 KO mice were lower than in WT mice in response to secondary infection (Fig. 8B). The differences between the antibody treatment and KO mice could be explained by an incomplete neutralization of IL-12p70 by the antibody *in vivo.* Nevertheless, both approaches showed that IL-12p70 was essential for increased NK-cell numbers and IFNγ production during secondary *T. gondii* infection.

In contrast to a complete absence of NK-cell IFNγ production after anti-IL-12p40 blockade, some percentage of NK cells still produced IFNγ in mice treated with anti-IL-12p70 and in IL-12p35 KO mice. One potential explanation for this is that there is an IL-12p70−independent and p40-dependent mechanism involved in NK-cell activation and IFNγ production during secondary infection. Because IL-12p40 is also a subunit of the cytokine IL-23, we tested whether IL-23 could also activate NK cells during secondary *T. gondii* infection. IL-23 extends the life of p40 KO mice during acute *T. gondii* infection (60). To find the role of IL-23 in activation of NK cells during reinfection, B6 mice were treated with anti-IL-23p19 5 to 6 wk after CPS immunization during secondary infection with RH. The number of NK cells was significantly lower in mice treated with anti-IL-23p19 as compared with control mice (Fig. 8C). The frequency of IFNγ+ NK cells after reinfection was not affected by IL-23p19 blockade. However, the absolute number of IFNγ-producing NK cells was reduced. These data show that during secondary infection IL-23 was essential for the increase in NK-cell numbers rather than IFNγ production. These data show that the NK-cell response to secondary *T. gondii* infection in the presence of immune memory is dependent on IL-12p70 and IL-23. IL-12p70 is essential for the NK-cell increase in numbers and for IFNγ production, and IL-23 is essential for the NK-cell increase in numbers.

### NK-cell maturation is reduced in the absence of IL-12 and IL-23

Based on the phenotypic analysis of NK cells responding to the secondary *T. gondii* infection, the majority (~73%) of IFNγ-producing NK cells expressed KLRG1 (Fig. 4D). KLRG1 expression is dependent on IL-12, which also induces expression of the transcription factor T-bet and IFNγ (61, 62). As IL-12p70 and IL-23 were required for NK-cell IFNγ production during secondary *T. gondii* infection, we next asked if these cytokines were also involved in the maturation of NK cells. The frequency of KLRG1+ NK cells was significantly lower in the absence of IL-12p40 and IL-23 after anti-IL-12p40 treatment and in the absence of IL-12p70 in IL-12p35 KO mice (Fig. 9A, 9B). In addition, the KLRG1+ NK cells were also reduced in the absence of IL-23 after anti-IL-23p19 treatment (Fig. 9C). Thus, NK cell maturation during secondary infection is also dependent on both IL-12p70 and IL-23.

**FIGURE 9.**
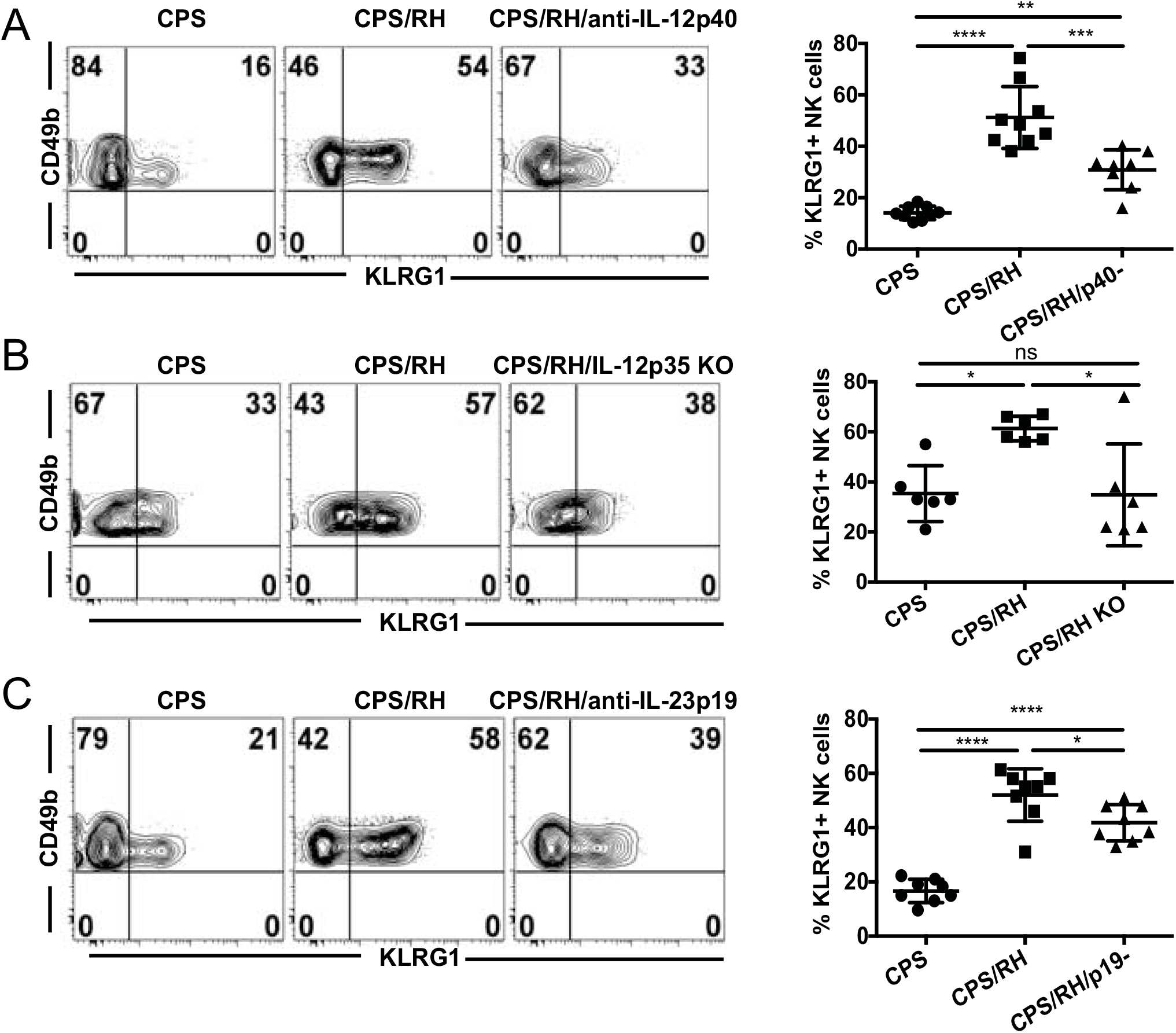
NK-cell maturation is reduced in the absence of IL-12 and IL-23 during secondary *T. gondii* infection. **(A-C)** B6 (**A** and **C**) and IL-12p35 KO **(B)** mice were i.p. infected with 1 × 10^6^ CPS1-1 and 5–6 wk later were i.p. infected with 1 × 10^3^ RH tachyzoites. KLRG1 expression on NK cells (CD49b+NKp46+CD3− live lymphocytes) was analyzed by flow cytometry in PECs at 3 d after RH infection. **(A)** Mice were treated with anti-IL-12p40 blocking antibody during RH infection. Representative contour plots and graphs from one of two independent experiments. Concatenated graphs from two independent experiments, n = 3–5 mice/group. **(B)** NK-cell KLRG1 expression in IL-12p35 KO mice. Representative contour plots from one of two independent experiments. Concatenated graphs from two independent experiments, n = 3 mice/group. **(C)** Mice were treated with anti-IL-23p19 antibody during RH infection. Representative plots and graphs from one of two independent experiments. Concatenated graphs from two independent experiments, n = 3 or 4 mice/group. Data are the mean ± SD. ns, not significant; *p < 0.05, **p < 0.01, ***p < 0.001, ****p < 0.0001, one-way ANOVA.

## Discussion

Recent evidence suggests that NK cells can participate in adaptive immunity by developing memory-like features or as immune effector cells regulated by memory T cells (3, 28, 31, 39, 51, 54). Whether NK cells contribute to long-term adaptive immune responses to *T. gondii* infection is not known. In this study we used a vaccine challenge model of *T. gondii* infection and showed that NK cells were required for protection during secondary infection when T-cell memory was present. However, adoptive transfer and fate mapping experiments indicated that NK cells did not develop cell-intrinsic memory-like characteristics and did not persist after primary *T. gondii* infection. NK-cell responses to secondary infection were dependent on new NK-cell generation and required cell-extrinsic factors IL-12/23p40, IL-12p70 and IL-23 cytokines to become activated and to become mature. Although NK cells did not develop memory-like features during *T. gondii* infection, they were essential for control of secondary parasite infection in an IL-12p70− and IL-23−dependent manner. Our results demonstrate a novel role for NK cells as innate immune cells that contribute significantly to secondary *T. gondii* infection, and both IL-12p70 and IL-23 are involved in this process. This study highlights the potential for NK cells to boost adaptive immunity against secondary infection in individuals whose immune systems become compromised.

Protection against secondary *T. gondii* infection is provided mainly by memory CD8+ T cells and was defined using vaccine and challenge approaches (35, 63, 64). How other immune cells, including NK cells, might contribute to this protection has not been clear. In a previous *T. gondii* vaccine challenge study performed in mice deficient for CD8+ T cells, cells expressing markers of NK cells including NK1.1 and ASGM1 provided protection against secondary infection (32). This study suggested that NK cells could substitute for CD8+ T cells in immunocompromised mice during an adaptive immune recall response. Whether this was also occurring in immunocompetent mice was not known. In our study, we demonstrated that NK1.1+ cells are critical for protection in immunocompetent mice in the presence of memory CD4+ and CD8+ T cells. Anti-NK1.1 treatment of vaccinated animals prior to lethal challenge resulted in significantly increased parasite burdens both in short-term and long-term infection. In addition, NK1.1+CD3− NK cells increased with a similar kinetic profile as compared with CD8+ T cells at the site of reinfection. Thus, we found that in addition to CD8+ T cells, NK cells contributed to control of secondary *T. gondii* infection in the presence of intact T-cell memory.

NK cells protect against primary *T. gondii* infection by producing IFNγ rather than by direct lysis of infected cells (16, 17, 26). We found that NK cells produce IFNγ, become cytotoxic (CD107a+) and are polyfunctional (IFNγ+CD107a+) in response to secondary infection. However, whereas the frequency of IFNγ-producing NK cells increased, the level of NK-cell cytotoxicity did not increase after reinfection. This suggests that IFNγ production could be the main NK-cell function necessary for protection during secondary parasite challenge in a manner similar to that which occurs during primary *T. gondii* infection (16–21).

A hallmark of NK cells that develop memory-like features is their ability to respond more efficiently and robustly to a secondary challenge similar to memory CD8+ T cells (2, 3, 28). For example, in a contact hypersensitivity model, hapten-sensitized *Rag2^-/-^* mice, which have neither T cells nor B cells, are more sensitive to secondary challenge than non-sensitized mice (2). Ly49H+ NK cells in mice infected with MCMV develop a more robust response to secondary MCMV infection (3). T and B cell−deficient mice immunized with influenza or HIV-1 virus-like particles (VLPs) have an increased response to secondary exposure to the corresponding VLPs (29). NK cells cultured with IL-12, IL-15 and IL-18 also develop the ability to respond more robustly to secondary stimulation (28). Whether primary *T. gondii* infection induces NK cells to develop a more efficient response to secondary parasite infection is not known. Our data indicate that *T. gondii* immunization of WT mice did not result in a more robust or efficient secondary NK-cell response during challenge infection. *T. gondii* immunization of T and B cell−deficient *(Rag1^-/-^)* mice did not lead to the development of stronger NK cell−dependent protection against secondary parasite infection. Expression of RAG1 early during NK-cell differentiation in the bone marrow has been suggested to be required for development of memory-like features in response to MCMV (48). Therefore, this may be one reason why we did not observe better NK cell−dependent protection in the *Rag1*^-/-^ mice. However, there was no increase in protection against parasite challenge in immunodeficient *Rag2^-/-^γc^-/-^* or WT mice that received purified *T. gondii*–experienced NK cells from immunized WT mice. Thus, our data suggest that although NK cells are an important component for the control of *T. gondii* challenge infection in the presence of T-cell memory, they do not appear to acquire cell-intrinsic features of a memory-like phenotype.

NK cells that develop memory-like features also acquire the ability to persist long-term(3, 31). They can also have specific tissue tropism (2, 3, 31). For example, liver NK cells develop a hapten-specific response (2). Liver and lung NK cells from influenza-immunized and liver NK cells from attenuated VSV-infected *Rag1*^-/-^ donors mediate protection in *Rag2^-/-^γc^-/-^* recipients (29). MCMV- and SIV-specific NK cells are found in the spleen and other organs (3, 31). Our fate mapping studies using reporter NKp46-CreERT2 × Rosa26-YFP mice indicate that, similar to their protective qualities, *T. gondii*–experienced and naïve NK cells were not intrinsically different in their ability to persist, in contrast to the MCMV-mediated increase in the number of long-lived YFP+ NK cells (9). We also did not observe any increase in YFP+ NK cells after *T. gondii* immunization in the peritoneum, spleen or bone marrow. This further supported the finding that NK cells do not develop cell-intrinsic characteristics of immune memory in response to *T. gondii* infection. Our results raise an important question: Why do NK cells not further differentiate and acquire memory-like characteristics during *T. gondii* infection as compared with hapten, virus or cytokine stimulation? Answering this question could explain why this parasite can escape and persist and be a continual problem for the host and also could help improve vaccine design.

NK-cell memory might not be generated in response to *T. gondii* infection for several reasons. These include the nature of NK-cell recognition of *T. gondii* infection or the lack thereof, co-stimulatory factors and the cytokine milieu. The development of memory-like features in NK cells in response to MCMV and HCMV infections requires signal 1 (antigen), signal 2 (co-stimulation) and signal 3 (cytokines) (3, 41, 43, 65, 66). One explanation for NK cells not developing intrinsic memory to *T. gondii* infection could be that the parasite does not itself express or does not induce a host cell to express a specific activating ligand for NK cells, and thus activation of NK cells lacks signal 1. In support of this possibility, our data indicate that after *T. gondii* immunization or after immunization and challenge no dominant NK-cell receptor was enriched within the responding population of total NK cells. We also do not observe any specific NK−cell receptor enrichment among the total responding NK-cell population during acute *T. gondii* infection, regardless of parasite virulence (42). Overall, we observed a wide array of NK-cell populations based on NK-cell receptor expression, indicating that there was a global response to infection. This could be common among apicomplexan infections because the NK-cell response to *Plasmodium falciparum* infection is also cell extrinsic (53). It is possible that NK cells evolved specific recognition mechanisms for viruses and bacteria but not for protozoan pathogens.

NK cells might not develop a memory-like response to *T. gondii* because they do not receive the necessary co-stimulatory signals or combination of cytokine stimulation. In response to MCMV and HCMV, specific co-stimultatory molecules and their interactions with their
 cognate ligands are essential for the generation of memory NK cells (43, 66, 67). DNAM-1 is important for memory NK-cell development during MCMV infection (43). We found that DNAM-1 co-stimulation was not required for the NK-cell response because its blockade did not alter NK-cell expansion or IFNγ production during secondary *T. gondii* infection. Other costimulatory molecules could also be involved. CD28 is important for maximizing the NK-cell responses to acute *T. gondii* infection (68). In addition to a potential alteration of co-stimualtory signals, *T. gondii* could induce higher expression of co-inhibitory molecules TIGIT and CD96 (69), which could then inhibit NK-cell memory formation; however, we did not observe any significant increases in the expression of these molecules during secondary *T. gondii* infection (data not shown).

Type I IFN and IL-12 are essential for expansion and survival of MCMV-specific NK cells (65, 70). Cytokine-induced memory-like NK cells can be generated in mice and humans after stimulation with IL-12, IL-18 and IL-15 (6, 28). All of these cytokines are produced during acute *T. gondii* infection (17, 42, 71, 72), with the exception of Type I IFNs (73). Therefore, it is surprising that NK cells do not further differentiate into memory-like cells. The development of NK-cell memory-like responses may require an activating receptor, costimulatory molecules and the correct combination of cytokines during *T. gondii* infection. Dissecting the reasons why NK cells do not develop memory-like features to *T. gondii* in a cell-intrinsic manner will be important future questions to address.

Because a second wave of NK-cell responses was critical for the control of secondary *T. gondii* infection in the presence of memory T cells, we addressed how the NK-cell extrinsic immune environment of vaccinated animals regulated their response. After *Rabies* virus vaccination secondary NK-cell responses depend on the presence of memory T cells (51). During *Rabies* virus secondary challenge, IL-2 producing antigen-specific memory CD4+ T cells in combination with IL-12 and IL-18 from accessory cells reactivated NK-cells. NK-cell responses in PBMCs of patients vaccinated with a *Plasmodium falciparum* specific vaccine correlated with IL-2 production by T cells (54). These studies suggest that memory T cells could provide cell extrinsic signals important for secondary NK cell responses to these infections. Unlike these previous studies, we did not find a role for antigen-specific memory CD4+ or CD8+ T cells in helping NK-cell responses to secondary infection. Activation of NK cells that is independent of memory T cells may have important implications for individuals that are T-cell deficient.

Multiple studies demonstrate that IL-12p40 is essential for protective immunity against *T. gondii* infection, including for NK cell IFNγ production during primary infection (17, 18, 74–76). Our data indicate that NK-cell activation in the presence of memory T cells during secondary infection of immunized animals was also dependent on the IL-12 subunit p40. IL-12p40 is a subunit of bioactive IL-12p70 (IL-12) and also of IL-23 (77). IL-12 and IL-23 are both capable of inducing IFNγ production by NK and T cells (77–79). During primary *T. gondii* infection, IL-12 is the main mediator of protection, and IL-23 provides protection if IL-12 is absent (60). Surprisingly, that study also demonstrated that IL-12p35 KO mice survived longer than IL-12/23p40 KO mice during *T. gondii* infection. The difference in protection was independent of T cells, raising an interesting question about the role of each of these cytokines in activating the non−T cell compartment to control primary *T. gondii* infection. Our results demonstrated that both IL-12 and IL-23 contribute to NK-cell responses during secondary *T. gondii* infection. NK-cell numbers were reduced in the absence of IL-12 or IL-23 during secondary *T. gondii* infection. However, NK-cell IFNγ was only partially reduced in the absence of IL-12, and depletion of IL-23 did not affect the frequency of IFNγ+ NK cells at the site of infection. Using anti-IL-12p40 treatment, we observed a near complete elimination of the secondary NK-cell response during parasite challenge. Thus, IL-12 is important in activating NK cells to produce IFNγ, whereas IL-23 may be more important for increasing the number of NK cells at the site of secondary infection in the presence of memory T cells. The mechanisms by which IL-12 and IL-23 work together to regulate NK-cell numbers and their IFNγ production in a *T. gondii* vaccine challenge situation could be via inducing NK-cell proliferation, migration and the ability to survive after infection.

In summary, we demonstrated that NK cells play an important protective role beyond primary *T. gondii* infection and in the presence of memory T cells after vaccination and during a challenge infection. NK cells become activated and help control the parasite during secondary infection. In their absence, parasite burdens are increased both in short-term and long-term challenge infection. NK cells do not develop cell-intrinsic characteristics of memory and rely on cell-extrinsic IL-12 and IL-23 to respond during secondary challenge. Thus, NK cells provide protection in the presence of memory T cells and should be taken into consideration in vaccination and immunotherapy strategies to combat *T. gondii* infection.

## Abbreviations used in this article

BM: bone marrow
HCMV: human cytomegalovirus
i.g.: intragastrically
MCMV: murine cytomegalovirus

