## Supplemental Figure 1 for "The IL-12– and IL-23–Dependent NK-Cell Response is Essential for Protective Immunity Against Secondary *Toxoplasma Gondii* Infection"

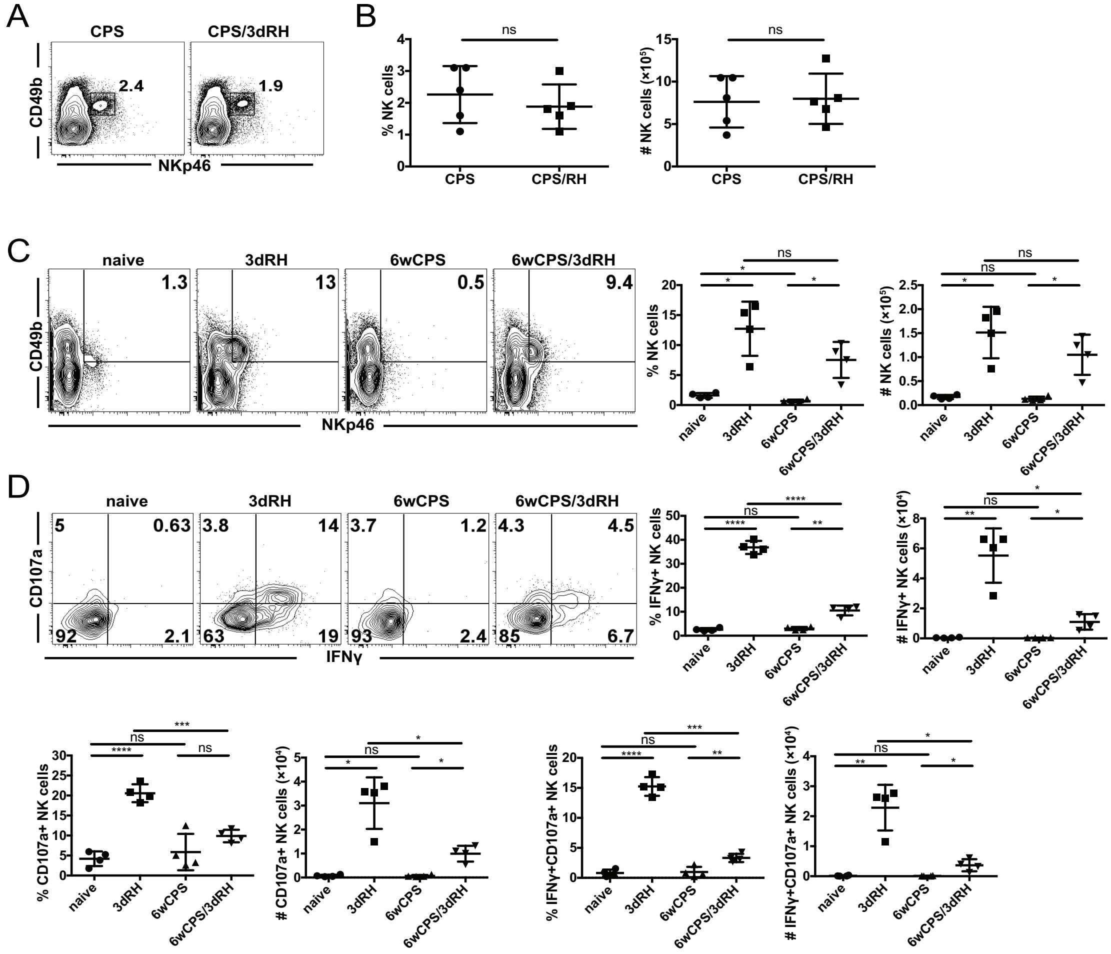


**Supplemental Figure 1.** NK cells become activated during an adaptive recall response. (**A, B**) B6 mice were infected i.p. with 1 × $10^6$ CPS and then infected i.p. with 1 × $10^3$ RH tachyzoites 5−6 wk later. Spleens were analyzed by flow cytometry 3 d after RH infection. (**A, B**) The percent and (**B**) number of NK cells (CD49b+NKp46+ in CD3− live lymphocytes) per spleen. Representative data from one of six independent experiments are shown, n = 3−5 mice/group. (**C, D**) PECs were harvested from naïve B6 mice and from B6 mice 3 d after RH infection (1 × $10^3$ RH tach.; 3dRH), 6 wk after CPS infection (1 × $10^6$ CPS tachyzoites; 6w CPS) and 3 d after RH reinfection (1 × $10^3$ RH tach.; 6w CPS/3dRH). (**C**) The percentage and number of NK cells per total PEC. (**D**) The percentage and number of IFNγ+, CD107a+ and IFNγ+CD107a+ NK cells per PEC. Data are representative of one of two independent experiments, n = 3 or 4 mice/group. Data are the mean ± SD. Unpaired Student's t-test with Welch's correction. ns, not significant; *p < 0.05, **p < 0.01, ***p < 0.001, ****p < 0.0001.

**Supplemental Figure 2.**

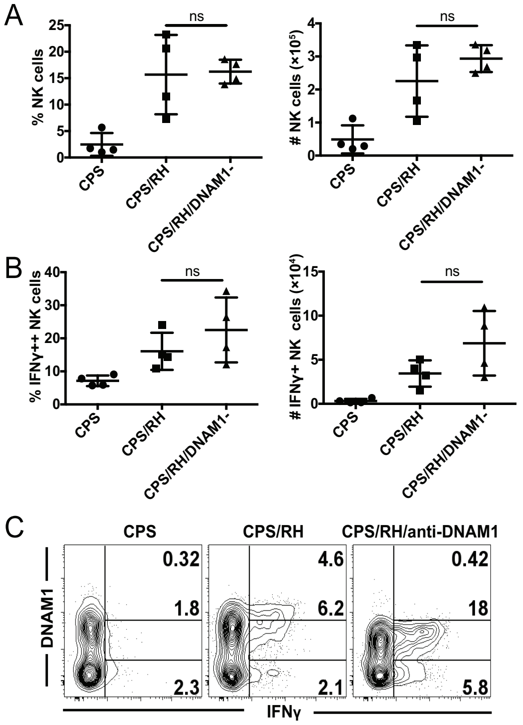


**Supplemental Figure 2.** NK cells become activated independent of DNAM-1 during secondary *T. gondii* infection. (**A**-**C**) B6 mice were infected i.p. with $1 \times 10^6$ CPS, reinfected i.p. with $1 \times 10^3$ RH tachyzoites 6 wk later and were treated with 100 µg i.p. anti-DNAM-1 on d −1 and 0. PECs were analyzed by flow cytometry at d 3 after RH infection. (**A**) Representative contour plots of NK cell (CD49b+NKp46+CD3− live lymphocytes) expression of DNAM-1 *vs.* IFNγ production. (**B** and **C**) The frequency and number of total **(B)** and IFNγ+ **(C)** NK cells. Data are the mean ± SD, n = 4 mice/group. ns, not significant, one-way ANOVA.

**Supplemental Figure 3.**

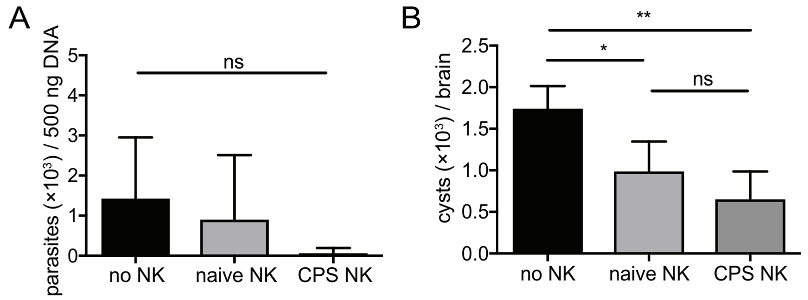


**Supplemental Figure 3.** *T. gondii*–experienced and naïve NK cells are not intrinsically different in their ability to protect. (**A** and **B**) NK cells were purified from spleens of naïve B6 mice (naïve NK) and from B6 mice 5 wk after CPS immunization (CPS NK) and i.v. transferred into B6 mice ($3 \times 10^6$ NK cells/mouse). (**A**) Parasite burdens were measured in recipient spleens by real-time PCR for the B1 *T. gondii* gene at 4 d after infection with $1 \times 10^5$ RH tachyzoites i.p. The data are from one experiment, n = 5 mice/group. (**B**) Cysts were counted in brains five weeks after infection with 10 ME49 cysts i.p. The data are from one experiment, n = 4 mice/group. ns, not significant; *p < 0.05, **p < 0.01, one-way ANOVA.

**Supplemental Figure 4.**

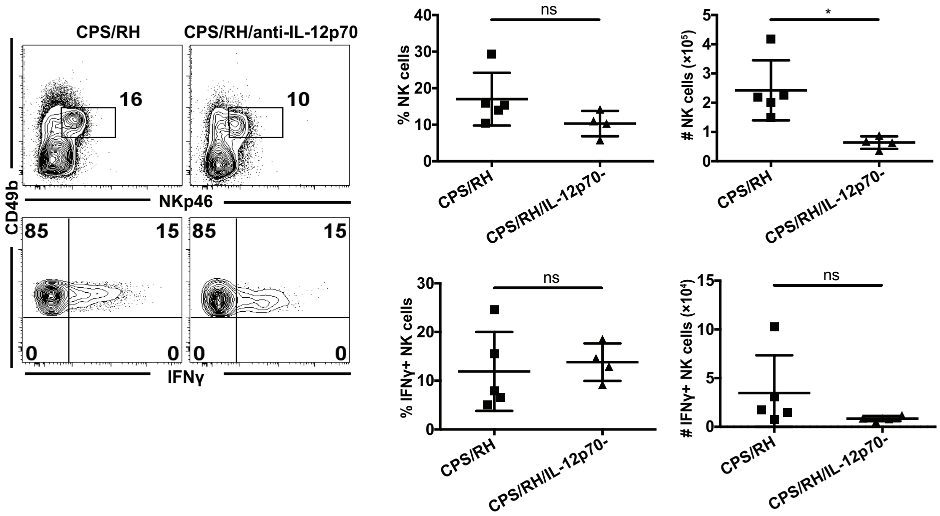


**Supplemental Figure 4.** IL-12p70 is required for the NK-cell response to secondary *T. gondii* infection. B6 mice were i.p. infected with $1 \times 10^6$ CPS1-1 and 6 wk later were treated with IL-12p70 neutralizing antibody or were untreated during i.p. infection with $1 \times 10^3$ RH tachyzoites. NK cells (CD49b+NKp46+CD3− lymphocytes) and their IFNγ production were analyzed in PECs at 3 d after RH infection by flow cytometry. Representative contour plots and graphs from one of two independent experiments, n = 4 or 5 mice/group. Data are the mean ± SD. ns, not significant; $*p < 0.05$, unpaired Student's t-test with Welch's correction.
